# Prediction tendency, eye movements, and attention in a unified framework of neural speech tracking

**DOI:** 10.1101/2023.06.27.546746

**Authors:** Juliane Schubert, Quirin Gehmacher, Fabian Schmidt, Thomas Hartmann, Nathan Weisz

**Affiliations:** Paris-Lodron-University of Salzburg, Department of Psychology, Centre for Cognitive Neuroscience, Salzburg, Austria; Neuroscience Institute, Christian Doppler University Hospital, Paracelsus Medical University, Salzburg, Austria; Department of Experimental Psychology, University College London, United Kingdom; Wellcome Centre for Human Neuroimaging, University College London, United Kingdom

**Author notes:** Address for correspondence Juliane Schubert, University of Salzburg, Center for Cognitive Neuroscience Hellbrunnerstr. 34, A-5020 SALZBURG, Austria/Europe. contributed equally (Juliane Schubert, Quirin Gehmacher).

**Keywords:** *active auditory sensing*, *speech processing*, *predictive processing*, *selective attention*, *eye movements*, *magnetoencephalography*

## Abstract

Auditory speech comprehension is a multi-faceted process in which attention, prediction, and sensorimotor integration (via active sensing) interact with or complement each other. Although different conceptual models that focus on one of these aspects exist, we still lack a unified understanding of their role in speech processing. Here, we first replicated two recently published studies from our lab, confirming 1) a positive relationship between individual prediction tendencies and neural speech tracking, and 2) the phenomenon of ocular speech tracking - the tracking of attended speech by eye movements - and its shared contribution with neural activity to speech processing. In addition, we extended these findings with complementary analyses and investigated these phenomena in relation to each other in a multi-speaker paradigm with continuous, narrative speech. Importantly, prediction tendency and ocular speech tracking seem to be unrelated. In contrast to the shared contributions of oculomotor and neural activity to speech processing over a distributed set of brain regions that are critical for attention, individual prediction tendency and its relation to neural speech tracking seem to be largely independent of attention. Based on these findings, we propose a framework that aims to bridge the gaps between attention, prediction, and active (ocular) sensing in order to contribute to a holistic understanding of neural speech processing. In this speculative framework for listening, auditory inflow is, on a basic level, temporally modulated via active ocular sensing, and incoming information is interpreted based on probabilistic assumptions.

## INTRODUCTION

Listening challenges the brain to infer meaning from vastly overlapping spectrotemporal information. For example, in complex, everyday environments, we need to select and segregate streams of speech from different speakers for further processing. To accomplish this task, (mainly independent) research lines suggested contributions of predictive and attentional processes to speech perception.

Predictive brain accounts (K. Friston, 2010; K. J. Friston et al., 2021; Knill & Pouget, 2004; Yon et al., 2019) suggest an active engagement in speech perception. In this view, experience-based internal models constantly generate and continuously compare top-down predictions with bottom-up input, thus inferring sound sources from neural activity patterns. This idea is supported by the influential role of speech predictability (e.g. semantic context and word surprisal) on speech processing in naturalistic contexts (Broderick et al., 2019; Donhauser & Baillet, 2020; Weissbart et al., 2020).

Selective attention describes the process by which the brain prioritises information to focus our limited cognitive capacities on relevant inputs while ignoring distracting, irrelevant ones. Its beneficial role in the processing of complex acoustic scenes, like the cocktail-party scenario (Zion Golumbic et al., 2013; Zion-Golumbic & Schroeder, 2012), supports the key role of attentional and predictive processes in natural listening. It is important to note that even “normal hearing” individuals vary greatly in everyday speech comprehension and communication (Ruggles et al., 2011), which consequently promotes interindividual variability in predictive processes (Siegelman & Frost, 2015) or selective attention (Oberfeld & Klöckner-Nowotny, 2016) as underlying modulators.

In a recent study (Schubert et al., 2023), we quantified individual differences in predictive processes as the tendency to anticipate low-level acoustic features (i.e. prediction tendency). We established a positive relationship of this “trait” with speech processing, demonstrating that “neural speech tracking” is enhanced in individuals with stronger prediction tendency, independent of situative demands on selective attention. Furthermore, we found an increased tracking of words of high surprisal, demonstrating the importance of predictive processes in speech perception. Here (and in the following) “neural speech tracking” refers to a correlation coefficient between actual brain responses and responses predicted from an encoding model based solely on the speech envelope.

Another important aspect with regard to speech processing - that has been vastly overlooked so far in neuroscience - is active auditory sensing. The potential benefit of active sensing - the engagement of (covert) motor routines in acquiring sensory inputs - was outlined for other sensory modalities (Schroeder et al., 2010). In the auditory domain, and especially for speech perception, motor contributions to an increased weighting of temporal precision were suggested to modulate auditory processing gain in support of (speech) segmentation (Morillon et al., 2015). Along these lines, It has been shown that covert, mostly blink related eye activity aligns with higher-order syntactic structures of temporally predictable, artificial speech (i.e. monosyllabic words; Jin et al, 2018). In support of ideas that the motor system is actively engaged in speech perception (Limerman et al., 19985; Galantucci et al., 2006), the authors suggest a global entrainment across sensory and (oculo)motor areas which implements temporal attention.

In another recent study from our lab (Gehmacher et al., 2024), we showed that eye movements continuously track intensity fluctuations of attended natural speech, a phenomenon we termed ocular speech tracking. In the present study, we focused on gaze patterns rather than blink-related activity, both to replicate findings from Gehmacher et al. (2024) and because blink activity is unsuitable for TRF analysis due to its discrete and artifact-prone nature. Hence, “Ocular speech tracking” (similarly to “neural speech tracking”) refers to the correlation coefficient between actual eye movements and movements predicted from an encoding model based on the speech envelope.

We further established links to increased intelligibility with stronger ocular tracking, and provided evidence that eye movements and neural activity share contributions to speech tracking. Taking into account the aforementioned study by Schubert and colleagues (2023), the two recently uncovered predictors of neural tracking (individual prediction tendency and ocular tracking) raise several empirical questions regarding the relationship between predictive processes, selective attention, and active ocular sensing in speech processing:

1. Are predictive processes related to active ocular sensing in the same way they are to neural speech tracking? Specifically, do individuals with a stronger tendency to anticipate predictable auditory features, as quantified through pre-stimulus neural representations in an independent tone paradigm, show increased or even decreased ocular speech tracking, measured as the correlation between predicted and actual eye movements? Or is there no relationship at all?
2. To what extent does selective attention influence the relationship between prediction tendency, neural speech tracking, and ocular speech tracking? For example, does the effect of prediction tendency or ocular speech tracking on neural tracking differ between a single-speaker and multi-speaker listening condition?
3. Are individual prediction tendency and ocular speech tracking related to behavioral outcomes, such as comprehension and perceived task difficulty? Speech comprehension is assessed through accuracy on comprehension questions, and task difficulty is measured through subjective ratings.

Although predictive processes, selective attention, and active sensing have been shown to contribute to successful listening, their potential interactions and specific roles in naturalistic speech perception remain unclear. Addressing these questions will help disentangle their contributions and establish an integrated framework for understanding how neural and ocular speech tracking support speech processing.

Here, we set out to answer these questions by a) replicating aforementioned key findings in a single experiment, and b) operationalising prediction tendency, active ocular sensing, and attention in an integrated framework of neural speech tracking. The purpose of this conceptual framework will be to organise these entities according to their assumed function for speech processing and to describe their relationship with each other. We therefore repeated the study protocol of Schubert et al. (2023) with slight modifications: Again, participants performed a separate, passive listening paradigm (also see Demarchi et al., 2019) that allowed us to quantify individual prediction tendency. Afterward, they listened to sequences of audiobooks (using the same stimulus material), either in a clear speech (0 distractors) or a multi-speaker (1 distractor) condition. Simultaneously recorded magnetoencephalographic (MEG) and Eye Tracking data confirmed the previous findings of Schubert et al. (2023) and Gehmacher et al. (2024): 1) Individuals with a stronger prediction tendency showed an increased neural speech tracking over left frontal areas, 2) eye movements track acoustic speech in selective attention, and 3) further mediate neural speech tracking effects over widespread, but mostly auditory regions. Additionally, we found an increased neural tracking of semantic violations (compared to their lexically identical controls), indicating that surprisal evoked responses indeed encode information about the stimulus. Interestingly, we could not find this difference in semantic processing for ocular speech tracking. Finally, we behaviorally assessed speech comprehension by probing participants on story content. Responses indicate that weaker performance in comprehension was related to increased ocular speech tracking while we did not find a significant relation to neural speech tracking. The findings suggest a differential role of prediction tendency, eye movements, and attention in speech processing. Behavioural responses further indicate substantial differences in ocular and neural engagement and perceptual outcomes. Based on these findings, we propose a framework of neural speech tracking where anticipatory predictions support the interpretation of auditory input along the perceptual hierarchy while active ocular sensing increases the temporal precision of peripheral auditory responses to facilitate bottom-up processing of selectively attended input.

## METHODS

### Subjects

In total, 39 subjects were recruited to participate in the experiment. For 3 participants, eye tracker calibration failed and they had to abort the experiment. We further controlled for excessive blinking (> 50% of data samples) and eye movements away from fixation cross (> 50% of data samples exceeded ⅓ of screen size on the horizontal or vertical plane) that suggest a lack of commitment to the experiment instructions. 7 subjects had to be excluded according to these criteria. Thus, a final sample size of 29 subjects (12 female, 17 male; mean age = 25.70, range = 19 - 43) was used for the analysis of brain and gaze data. All participants reported normal hearing and had normal, or corrected to normal, vision. They gave written, informed consent and reported that they had no previous neurological or psychiatric disorders. The experimental procedure was approved by the ethics committee of the University of Salzburg and was carried out in accordance with the declaration of Helsinki. All participants received either a reimbursement of 10 € per hour or course credits.

### Experimental Procedure

The current study is a conceptual replication of a previous experiment (for details see Schubert et al., 2023), using the same experimental structure and the same auditory stimuli. The current design only differs in that one condition was dropped and the whole experiment was shortened (participants spent approximately 2 hours in the MEG including preparation time). In addition to the previous study, this time we recorded ocular movements (also see Data acquisition and Preprocessing).

Before the start of the experiment, participants’ head shapes were assessed using cardinal head points (nasion and pre-auricular points), digitised with a Polhemus Fastrak Digitiser (Polhemus), and around 300 points on the scalp. For every participant, MEG sessions started with a 5-minute resting-state recording, after which the individual hearing threshold was determined using a pure tone of 1043 Hz. This was followed by 2 blocks of passive listening to tone sequences of varying entropy levels to quantify individual prediction tendencies (see Quantification of individual prediction tendency and Figure 1A-C). Participants were instructed to look at a black fixation cross at the centre of a grey screen. In the main task, 4 different stories were presented in separate blocks in random order and with randomly balanced selection of the target speaker (male vs. female voice). Each block consisted of 2 trials with a continuous storyline, with each trial corresponding to one of 2 experimental conditions: a single speaker and a multi-speaker condition (see also Figure 1D). The distractor speaker was always of the opposite sex of the target speaker (and was identical to the target speaker in a different run). Distracting speech was presented exactly 20 s after target speaker onset, and all stimuli were presented binaurally at equal volume (40db above individual hearing threshold) for the left and right ear (i.e. at phantom centre). Participants were instructed to attend to the first speaker and their understanding was tested using comprehension questions (true vs. false statements) at the end of each trial (e.g.: “Das Haus, in dem Sofie lebt, ist rot” *(The house Sofie lives in is red)*, “Ein gutes Beispiel für unterschiedliche Dialekte sind die Inuit aus Alaska und Grönland” *(A good example of different dialects are the Inuit from Alaska and Greenland)*…). Furthermore, participants indicated their task engagement and their perceived task-difficulty on a 5-point Likert scale at the end of every trial. During the audiobook presentation, participants were again instructed to look at the fixation-cross on the screen and to blink as little as (still comfortably) possible. The experiment was coded and conducted with the Psychtoolbox-3 (Brainard, 1997; Kleiner et al., 2007), with an additional class-based library (‘Objective Psychophysics Toolbox’, o_ptb) on top of it (Hartmann & Weisz, 2020).

**Figure 1:**
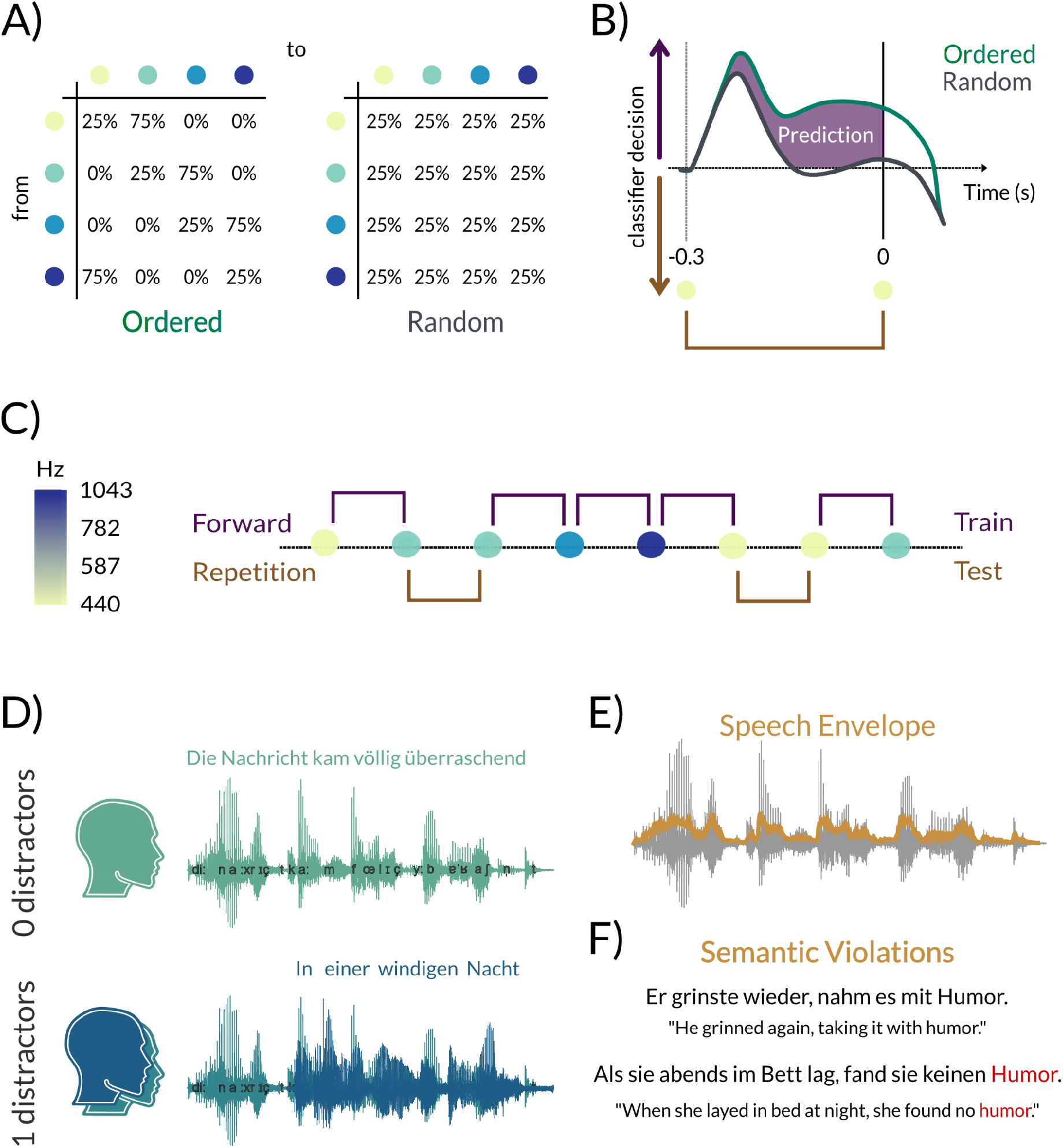
*Quantification of individual prediction tendency and the multi-speaker paradigm.* **A)** Participants passively listened to sequences of pure tones in different conditions of entropy (ordered vs. random). Four tones of different fundamental frequencies were presented with a fixed stimulation rate of 3 Hz, their transitional probabilities varied according to respective conditions. **B)** Expected classifier decision values contrasting the brains’ prestimulus tendency to predict a forward transition (ordered vs. random). The purple shaded area represents values that were considered as prediction tendency **C)** Exemplary excerpt of a tone sequence in the ordered condition. An LDA classifier was trained on forward transition trials of the ordered condition (75% probability) and tested on all repetition trials to decode sound frequency from brain activity across time. **D)** Participants either attended to a story in clear speech, i.e. 0 distractor condition, or to a target speaker with a simultaneously presented distractor (blue), i.e. 1 distractor condition. **E)** The speech envelope was used to estimate neural and ocular speech tracking in respective conditions with temporal response functions (TRF). **F)** The last noun of some sentences was replaced randomly with an improbable candidate to measure the effect of envelope encoding on the processing of semantic violations. Adapted from Schubert et al., 2023

### Stimuli

We used the same stimulus material as in Schubert et al. (2023), recorded with a t.bone SC 400 studio microphone at a sampling rate of 44100 Hz. In total, we used material from 4 different, consistent stories (see Table 1). These stories were split into 3 separate trials of approximately 3 - 4 min. The first two parts of each story were always narrated by a target speaker, whereas the last part served as distractor material. Additionally, we randomly selected half of the nouns that ended a sentence (N = 79) and replaced them with the other half to induce unexpected semantic violations. The swap of nouns happened in the written script before the audio material was recorded in order to avoid any effects of audio clipping. Narrators were aware of the semantic violations and had been instructed to read out the words as normal. Consequently all target words occurred twice in the text, once in a natural context (serving as “lexical controls”) and once in a mismatched context (serving as “semantic violations”) within each trial, resulting in two sets of lexically identical words that differed greatly in their contextual probabilities (see Figure 1F for an example). Participants were unaware of these semantic violations. All trials were recorded twice, narrated by a different speaker (male vs. female). Stimuli were presented in 4 blocks containing 2 trials each (a single and a multi-speaker trial), resulting in 2 male and 2 female target speaker blocks for every participant.\

**Table 1:**
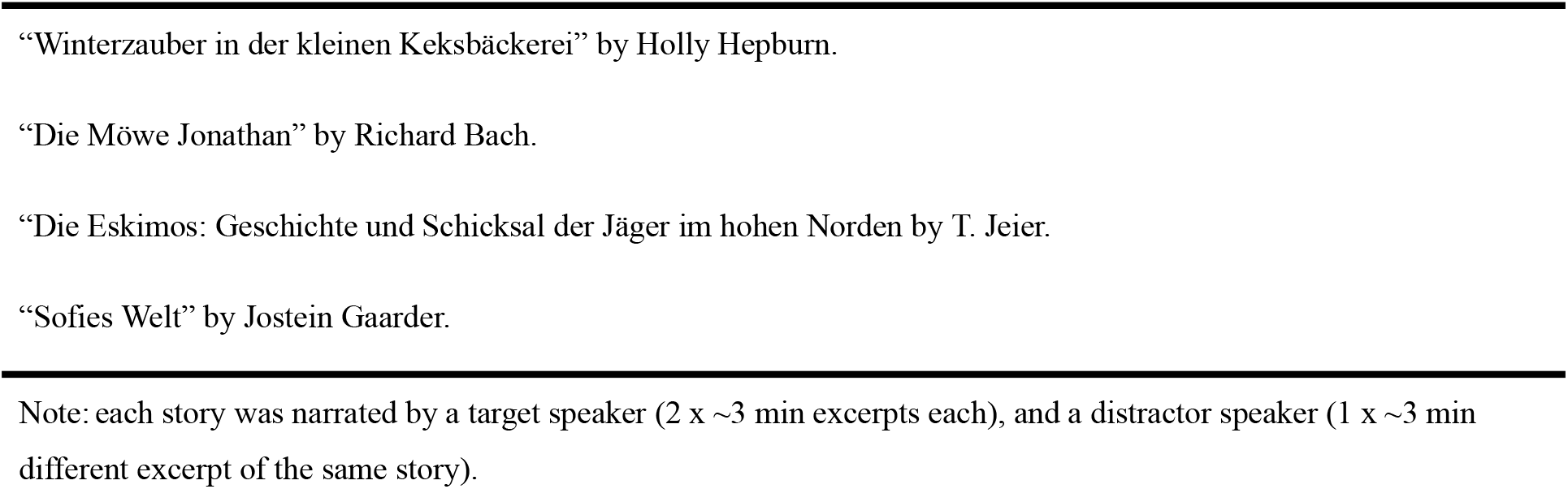
List of all ebooks and short stories that were used as a basis for audio material Note: each story was narrated by a target speaker (2 x ∼3 min excerpts each), and a distractor speaker (1 x ∼3 min different excerpt of the same story).

### Data Acquisition and Preprocessing

Brain activity was recorded using a whole head MEG system (Elekta Neuromag Triux, Elekta Oy, Finland), placed within a standard passive magnetically shielded room (AK3b, Vacuumschmelze, Germany). We used a sampling frequency of 1 kHz (hardware filters: 0.1 - 330 Hz). The signal was recorded with 102 magnetometers and 204 orthogonally placed planar gradiometers at 102 different positions. In a first step, a signal space separation algorithm, implemented in the Maxfilter program (version 2.2.15) provided by the MEG manufacturer, was used to clean the data from external noise and realign data from different blocks to a common standard head position. Data preprocessing was performed using Matlab R2020b (The MathWorks, Natick, Massachusetts, USA) and the FieldTrip Toolbox (Oostenveld et al., 2011). All data was filtered between 0.1 Hz and 30 Hz (Kaiser windowed finite impulse response filter) and downsampled to 100 Hz. To identify eye-blinks and heart rate artefacts, 50 independent components were identified from filtered (0.1 - 100 Hz), downsampled (1000 Hz) continuous data of the recordings from the entropy modulation paradigm, and on average 3 components were removed for every subject. All data was filtered between 0.1 Hz and 30 Hz (Kaiser windowed finite impulse response filter) and downsampled to 100 Hz. Data of the entropy modulation paradigm was epoched into segments of 1200 ms (from 400 ms before sound onset to 800 ms after onset). Multivariate pattern analysis (see quantification of individual prediction tendency) was carried out using the MVPA-Light package (Treder, 2020). Data of the listening task was temporally aligned with the corresponding speech envelope, which was extracted from the audio files using the Chimera toolbox (Smith et al., 2002) over a broadband frequency range of 100 Hz - 10 kHz (in 9 steps, equidistant on the tonotopic map of auditory cortex; see also Figure 1E).

Eye tracking data from both eyes were acquired using a Trackpixx3 binocular tracking system (Vpixx Technologies, Canada) at a sampling rate of 2 kHz with a 50 mm lens. Participants were seated in the MEG at a distance of 82 cm from the screen. Their chin rested on a chinrest to reduce head movements. Each experimental block started with a 13-point calibration / validation procedure that was used throughout the block. Blinks were automatically detected by the Trackpixx3 system and excluded from horizontal and vertical eye movement data. We additionally excluded 100 ms of data around blinks to control for additional blink artefacts that were not automatically detected by the eye tracker. We then averaged gaze position data from the left and right eye to increase the accuracy of gaze estimation (Cui & Hondzinski, 2006). Missing data due to blink removal were then interpolated with a piecewise cubic Hermite interpolation. Afterwards, data was imported into the FieldTrip Toolbox, bandpass filtered between 0.1 - 40 Hz (zero-phase finite impulse response (FIR) filter, order: 33000, hamming window), and cut to the length of respective speech segments of a block. Data were then downsampled to 1000 Hz to match the sampling frequency of neural and speech envelope data and further corrected for a 16 ms delay between trigger onset and actual stimulation. Finally, gaze data was temporally aligned with the corresponding speech envelope and downsampled to 100 Hz for TRF analysis after an antialiasing low-pass filter at 20 Hz was applied (zero-phase FIR filter, order: 662, hamming window).

### Quantification of Prediction Tendency

The quantification of individual prediction tendency was the same as in Schubert et al. (2023): We used an entropy modulation paradigm where participants passively listened to sequences of 4 different pure tones (f1: 440 Hz, f2: 587 Hz, f3: 782 Hz, f4: 1043 Hz, each lasting 100 ms) during two separate blocks, each consisting of 1500 tones presented with a temporally predictable rate of 3 Hz. Entropy levels (ordered / random) changed pseudorandomly every 500 trials within each block, always resulting in a total of 1500 trials per entropy condition. While in an “ordered” context certain transitions (hereinafter referred to as forward transitions, i.e. f1→f2, f2→f3, f3→f4, f4→f1) were to be expected with a high probability of 75%, self repetitions (e.g., f1→f1, f2→f2,…) were rather unlikely with a probability of 25%. However, in a “random” context all possible transitions (including forward transitions and self repetitions) were equally likely with a probability of 25% (see Figure 1A). This design gives us the opportunity to directly compare the processing of “predictable” and “unpredictable” sounds of the same frequency in a time-resolved manner.

To estimate the extent to which individuals anticipate auditory features (i.e. sound frequencies) according to their contextual probabilities, we used a multiclass linear discriminant analyser (LDA) to decode sound frequency (f1 - f4) from brain activity (using data from the 102 magnetometers) between -0.3 and 0 s. We specifically looked at a prestimulus window in order to capture top-down expectations driven by contextual regularity and temporal predictability in the design. The classification approach enables us to infer on “anticipatory predictions” (i.e. the representation of sound frequencies of high probabilities before their observation). Based on the resulting classifier decision values (i.e. d1 - d4 for every test-trial and time-point), we calculated **“**individual prediction tendency**”**. We define **“**individual prediction tendency**”** as the tendency to pre-activate sound frequencies of high probability (i.e. a forward transition from one stimulus to another: f1→f2, f2→f3, f3→f4, f4→f1). In order to capture any prediction-related neural activity, we trained the classifier exclusively on ordered forward trials (see Figure 1B). Afterwards, the classifier was tested on all self-repetition trials, providing classifier decision values for every possible sound frequency, which were then transformed into corresponding transitions (e.g. d1(t) | f1(t-1) “dval for 1 at trial t, given that 1 was presented at trial t-1” → repetition, d2(t) | f1(t-1) → forward,…). The tendency to represent a forward vs. repetition transition was contrasted for both ordered and random trials (see Figure 1C). Using self-repetition trials for testing, we ensured a fair comparison between the ordered and random contexts (with an equal number of trials and the same preceding bottom-up input). Thus, we quantified “prediction tendency” as the classifier’s pre-stimulus tendency to a forward transition in an ordered context exceeding the same tendency in a random context (which can be attributed to carry-over processing of the preceding stimulus). Then, using the summed difference across pre-stimulus times, one value can be extracted per subject (also see Figure 1B).

### Encoding Models

To quantify the neural representations corresponding to the acoustic envelope, we calculated a multivariate temporal response function (TRF) using the Eelbrain toolkit (Brodbeck et al., 2021). A deconvolution algorithm (boosting; David et al., 2007) was applied to the concatenated trials to estimate the optimal TRF to predict the brain response from the speech envelope, separately for each condition (single vs. multi-speaker). Before model fitting, MEG data of 102 magnetometers were normalised by subtracting the mean and dividing by the standard deviation (i.e. z-scoring) across all channels (as recommended by Crosse et al., 2016) Similarly, the speech envelope was also z-scored, however, after the transformation the negative of the minimum value (which naturally would be zero) was added to the time-series to retain zero values (z’ = z+(min(z)*-1). The defined time-lags to train the model were from -0.4 s to 0.8 s. To evaluate the model, the data was split into 4 folds, and a cross-validation approach was used to avoid overfitting (Ying, 2019). The resulting predicted channel responses (for all 102 magnetometers) were then correlated with the true channel responses to quantify the model fit and the degree of speech envelope tracking at a particular sensor location.

To investigate the effect of semantic violations, we used the same TRFs trained on the whole dataset (with 4-fold cross validation) since single word epochs would have been too short to derive meaningful TRFs. Instead the true as well as the predicted data was segmented into single word epochs of 2 seconds starting at word onset (using a forced-aligner; Kisler et al., 2017; Schiel & Ohala, 1999). We selected semantic violations as well as their lexically identical controls and correlated true with predicted responses for every word (thus, we conducted the same analysis as for the overall encoding effect, focusing on only part of the data). We then averaged the result within each condition (i.e. single vs. multi-speaker) and word type (i.e. high vs. low surprisal).

The same TRF approach was also used to estimate ocular speech tracking, separately predicting eye movements in the horizontal and vertical direction using the same time-lags (from -0.4 s to 0.8 s). The same z-scoring was applied to the speech envelope. However, horizontal and vertical eye channel responses were normalised within channels.

### Mediation Analysis

To investigate the contribution of eye movements to neural speech tracking, we approached a mediation analysis similar to Gehmacher et al. (2024). The TRFs that we obtained from these encoding models can be interpreted as time-resolved weights for a predictor variable that aims to explain a dependent variable (very similar to beta-coefficients in classic regression analyses). We simply compared the plain effect of the speech envelope on neural activity to its direct (residual) effect by including an indirect effect via eye movements into our model. In order to account for a reduction in speech envelope weights simply due to the inclusion of this additional (eye-movement) predictor, we obtained a control model by including a circularly shifted (by one-half of its total duration) version of the eye-movement predictor in addition to the unchanged speech envelope. Thus, the plain effect (i.e. speech envelope predicting neural responses) is represented in the absolute weights (i.e. TRFs) obtained from this control model with the speech envelope and time-shifted eye movement data as the predictor of neural activity. The direct (residual) effect (not mediated by eye movements) is obtained from the model including the speech envelope as well as true eye movements and is represented in the exclusive weights (c’) of the former predictor (i.e. speech envelope). If model weights are significantly reduced by the inclusion of true eye movements into the model in comparison to a model with a time-shuffled version of the same predictor (i.e. c’ < c), this indicates that a meaningful part of the relationship between the speech envelope and neural responses was mediated by eye movements (for further details see also Gehmacher et al., 2024).

### Source and Principal Component Analysis

In order to estimate the location along with the temporal profile of this mediation effect and at the same time minimise the number of comparisons, we computed the main components of the effect, projected into source space. For this, we used all 306 MEG channels for our models as described in the previous section (note that here MEG responses were z-scored within channel type, i.e. within magnetometers and gradiometers separately). The resulting envelope model weights were then projected into source-space using an LCMV beamforming approach (Van Veen et. al., 1997). Spatial filters were computed by warping anatomical template images to the individual head shape and further brought into a common space by co-registering them based on the three anatomical landmarks (nasion, left and right preauricular points) with a standard brain from the Montreal Neurological Institute (MNI, Montreal, Canada; Mattout et al., 2007). For each participant, a single-shell head model (Nolte, 2003) was computed. Finally, as a source model, a grid with 1 cm resolution and 2982 voxels based on an MNI template brain was morphed into the brain volume of each participant. This allowed group-level averaging and statistical analysis as all the grid points in the warped grid belong to the same region across subjects.

Afterwards, envelope TRF edges of both plain and direct models were cropped to time-lags from -0.3 - 0.7 s to exclude potential regression artefacts. We then subtracted the absolute direct from the absolute plain TRF to obtain the ‘abs’ indirect (mediation) effect. We transformed the 2982 voxel space into an orthogonal component space with a principal component analysis (PCA) based on the grand average mediation effect. The number of components for further analysis was visually determined by plotting the ranked cumulative explained variance and estimating the ‘elbow’. Based on this inspection, we extracted weight matrices of the first three components and multiplied individual ‘abs’ plain TRFs (from the control model with time-shuffled eye movements) and ‘abs’ direct TRFs (from the test model with true eye-movements) with this weight matrix to obtain individual source location and temporal profile of the mediation components. The projected, single subject data was then used for statistical analysis.

### Statistical Analysis and Bayesian Models

To calculate the statistics, we used Bayesian multilevel regression models with Bambi (Capretto et al., 2020), a python package built on top of the PyMC3 package (Salvatier et al., 2016), for probabilistic programming. First, as a conceptual replication of Schubert et al. (2023), we investigated the effect of neural speech tracking and its relation to individual prediction tendency as well as the influence of increasing noise (i.e. adding a distractor speaker). Separate models were calculated for all 102 magnetometers using the following model formula:

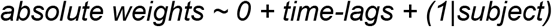

“Neural speech envelope tracking” refers to the correlation coefficients between predicted and true brain responses from the aforementioned encoding model, trained and tested on the whole audio material within condition (single vs. multi-speaker). (Note that prediction tendency was always z-scored before entering the models).

Similarly, we investigated the effect of ocular speech tracking under different conditions of attention (i.e. attended single speaker, attended multi-speaker and unattended multi-speaker). In addition to replicating the findings from Gehmacher et al. (2024), we extended this analysis for a detailed investigation of horizontal and vertical ocular speech envelope tracking and further included prediction tendency as a predictor:

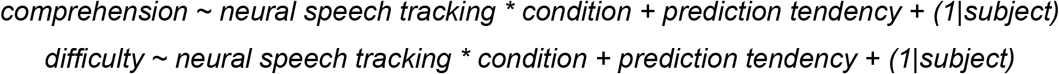

“Ocular speech envelope tracking” refers to the correlation coefficients between predicted and true eye movements from the encoding model (again trained and tested on the whole audio material within condition.

To investigate the effect of semantic violations, we compared envelope tracking between target words (high surprisal) and lexically matched controls (low surprisal), both for neural as well as ocular speech tracking:

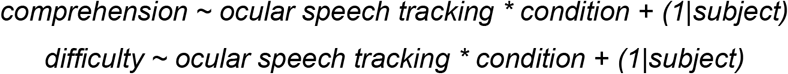

Here “Speech envelope tracking” refers to the correlation coefficients between predicted and true responses (either neural or ocular) from the encoding model trained on the whole audio material but tested solely on the word-segments.

For the mediation analysis, we compared weights (TRFs) for the speech envelope between a plain (control) model that included shuffled eye-movements as second predictor and a residual (test) model that included true eye movements as a second predictor (i.e. c’ < c). In order to investigate the temporal dynamics of the mediation, we included time-lags (-0.3 - 0.7 s) as a fixed effect into the model. To investigate potential null effects (neural speech tracking that is definitely independent of eye movements), we used a region of practical equivalence (ROPE) approach (see for example Kruschke, 2018). The dependent variable (absolute weights) was z-scored across time-lags and models to get standardised betas for the mediation effect (i.e. c’ < c):

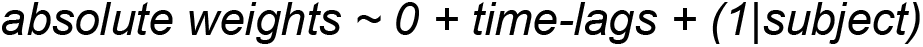

As suggested by Kruschke (2018), a null effect was considered for betas ranging between-0.1 and 0.1. Accordingly, it was considered a significant mediation effect if betas were above 0.1 and at minimum two neighbouring time-points also showed a significant result. Finally, we investigated the relationship between neural as well as ocular speech tracking and behavioural data using the averaged accuracy from the questions on story content that were asked at the end of each trial (in the following: “comprehension”) and averaged subjective ratings of difficulty. In order to avoid calculating separate models for all magnetometers again, we selected 10% of the channels that showed the strongest speech encoding effect and used the averaged speech tracking (z-scored within condition before entering the model) as predictor:

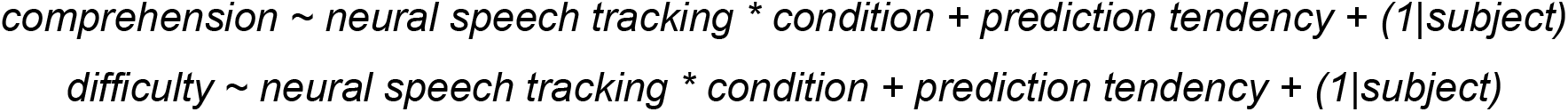

To investigate the relation between ocular speech tracking and behavioural performance, we used the following model (again speech tracking was z-scored within condition) separately for horizontal and vertical gaze direction:

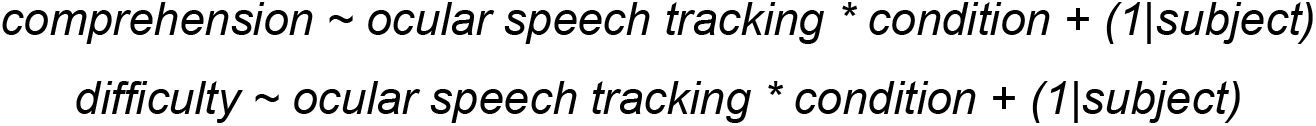

For all models (as in Gehmacher et al, 2024 and Schubert et al., 2023), we used the weakly-or non-informative default priors of Bambi (Capretto et al., 2020) and specified a more robust Student-T response distributions instead of the default gaussian distribution. To summarise model parameters, we report regression coefficients and the 94% high density intervals (HDI) of the posterior distribution (the default HDI in Bambi). Given the evidence provided by the data, the prior and the model assumptions, we can conclude from the HDIs that there is a 94% probability that a respective parameter falls within this interval. We considered effects as significantly different from zero if the 94%HDI did not include zero (with the exception of the mediation analysis where the 94%HDI had to fall above the ROPE). Furthermore, we ensured the absence of divergent transitions (r^ < 1.05 for all relevant parameters) and an effective sample size > 400 for all models (an exhaustive summary of Bayesian model diagnostics can be found in Vehtari et al., 2021). Finally, when we estimated an effect on brain sensor level (using all 102 magnetometers), we defined clusters for which an effect was only considered as significant if at minimum two neighbouring channels also showed a significant result.

## RESULTS

### Individual prediction tendency is related to neural speech tracking

In the first step, we wanted to investigate the relationship between individual prediction tendency and neural speech tracking under different conditions of noise. We quantified prediction tendency as the individual tendency to represent auditory features (i.e. pure tone frequency) of high probability in advance of an event (i.e. pure tone onset). Thus, this measure is a single value per subject, which comprises a) differences between two contextual probabilities (i.e. ordered vs. random) in b) feature-specific tone representations c) in advance of their observation (summed over a time-window of -0.3 - 0 s). Importantly, this prediction tendency was assessed in an independent entropy modulation paradigm (see Fig. 1). On a group level we found an increased tendency to pre-activate a stimulus of high probability (i.e. forward transition) in an ordered context compared to a random context (see Fig, 2A). This effect replicates results from our previous work (Schubert et al., 2023, 2024). Using the summed difference between entropy levels (ordered - random) across pre-stimulus time, one value was extracted per subject (Fig. 2B). This value was used as a proxy for “individual prediction tendency” and correlated with encoding of clear speech across different MEG sensors.

**Figure 2:**
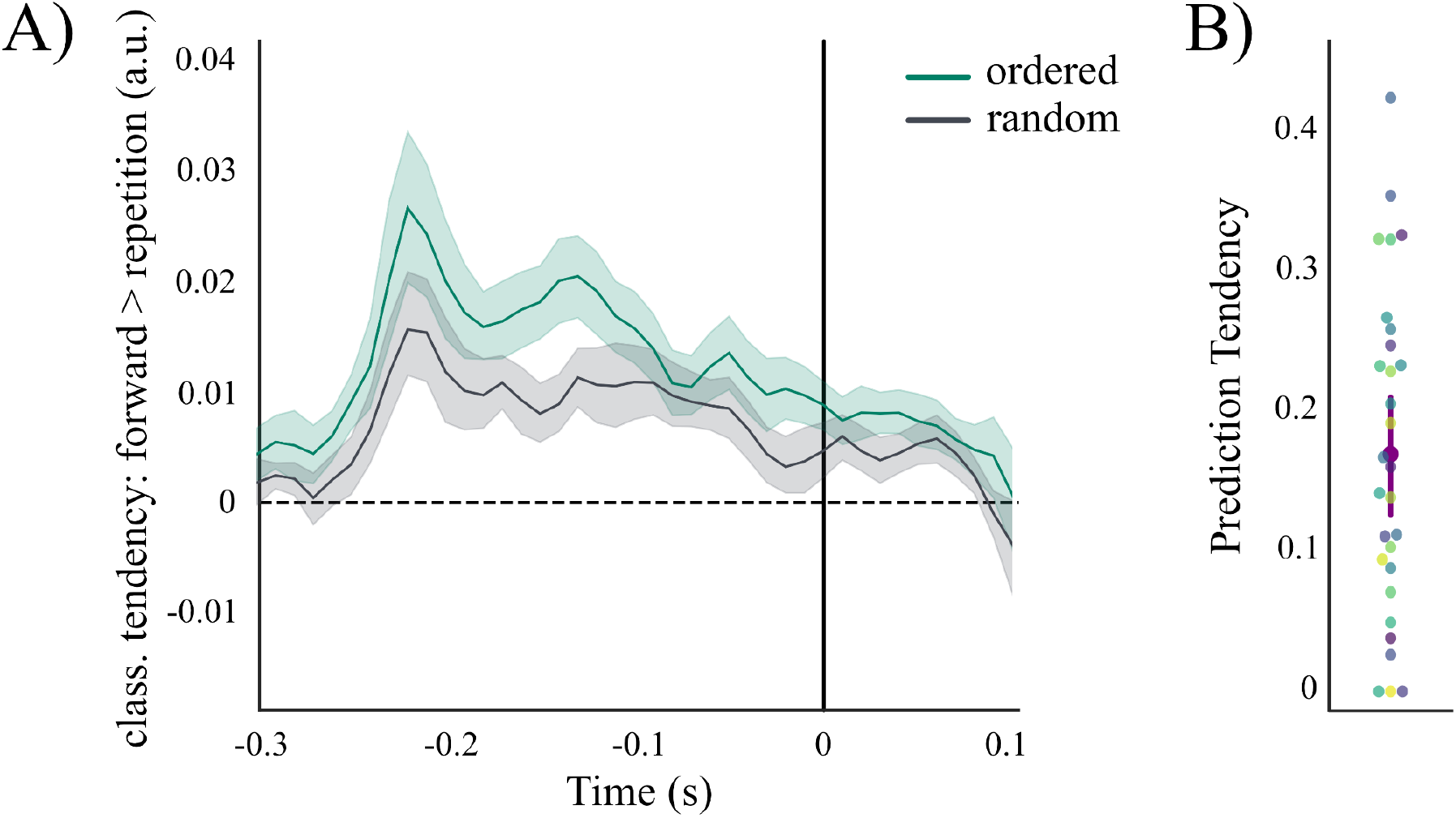
*Individual prediction tendency*. **A)** Time-resolved contrasted classifier decision: forward > repetition for ordered and random repetition trials. Classifier tendencies showing frequency-specific prediction for tones with the highest probability (forward transitions) can be found even before stimulus onset but only in an ordered context (shaded areas always indicate 95% confidence intervals). Using the summed difference across pre-stimulus time, one prediction value was extracted per individual subject. **B)** Distribution of prediction tendency values across subjects (N = 29).

Neural speech tracking, quantified as the correlation coefficients between predicted and observed MEG responses to the speech envelope, was used as the dependent variable in Bayesian regression models (Fig. 3A; see Fig. 3B for TRF weights). These models included condition (single vs. multi-speaker) as a fixed effect, with either prediction tendency or word surprisal as an additional predictor, and random effects for participants. Replicating previous findings (Schubert et al., 2023), we found widespread encoding of clear speech (average over cluster: *β* = 0.035, 94%HDI = [0.024, 0.046]), predominantly over auditory processing regions (Fig. 3C), that was decreased (*β* = -0.018, 94%HDI = [-0.029, -0.006]) in a multi-speaker condition (Fig. 3D). Furthermore, a stronger prediction tendency was associated with increased neural speech tracking (*β* = 0.014, 94%HDI = [0.004, 0.025]) over left frontal sensors (see Fig. 3E). We found no interaction between prediction tendency and condition (see Fig. 3F). These findings indicate that the relationship between individual prediction tendency and neural speech tracking is largely unaffected by demands on selective attention.

**Figure 3:**
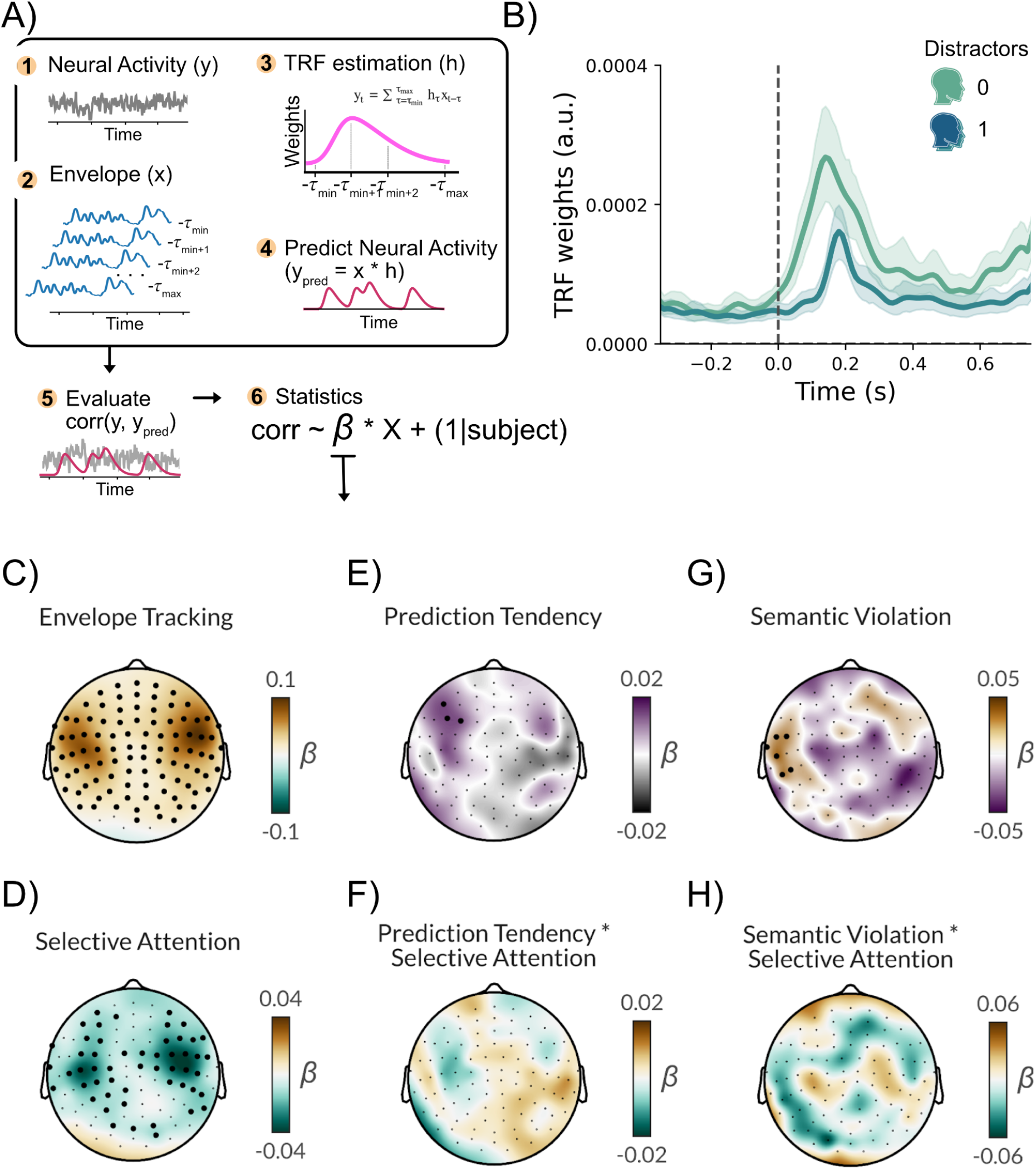
*Neural speech tracking is related to prediction tendency and word surprisal, independent of selective attention.* **A)** Envelope (x) - response (y) relationships are estimated using deconvolution (Boosting). The TRF (filter kernel, h) models how the brain processes the envelope over time. This filter is used to predict neural responses via convolution. Predicted responses are correlated with actual neural activity to evaluate model fit and the TRF’s ability to capture response dynamics. Correlation coefficients from these models are then used as dependent variables in Bayesian regression models. (Panel adapted from Gehmacher et al., 2024b). **B)** Temporal response functions (TRFs) depict the time-resolved neural tracking of the speech envelope for the single speaker and multi speaker target condition, shown here as absolute values averaged across channels. Solid lines represent the group average. Shaded areas represent 95% Confidence Intervals. **C–H)** The beta weights shown in the sensor plots are derived from Bayesian regression models in A. For Panel C, this statistical model is based on correlation coefficients computed from the TRF models (further details can be found in the Methods Section). **C)** In a single speaker condition, neural tracking of the speech envelope was significant for widespread areas, most pronounced over auditory processing regions. **D)** The condition effect indicates a decrease in neural speech tracking with increasing noise (1 distractor). **E)** Stronger prediction tendency was associated with increased neural speech tracking over left frontal areas. **F)** However, there was no interaction between prediction tendency and conditions of selective attention. **G)** Increased neural tracking of semantic violations was observed over left temporal areas. **H)** There was no interaction between word surprisal and speaker condition, suggesting a representation of surprising words independent of background noise. Marked sensors indicate ‘significant’ clusters, defined as at least two neighboring channels showing a significant result. N = 29.

Additionally, we wanted to investigate how semantic violations affect neural speech tracking. For this reason, we introduced rare words of high surprisal into the story by randomly replacing half of the nouns at the end of a sentence with the other half. In a direct comparison with lexically identical controls, we found an increased neural tracking of semantic violations (*β* = 0.039, 94%HDI = [0.007, 0.071]) over left temporal areas (see Fig. 3G). Furthermore, we found no interaction between word surprisal and speaker condition (see Fig. 3H). These findings indicate an increased representation of surprising words independent of background noise.

In sum, we found that individual prediction tendency as well as semantic predictability affect neural speech tracking.

### Eye movements track acoustic speech in selective attention

In a second step, we aimed to replicate previous findings from Gehmacher and colleagues (2024), showing that eye movements track the acoustic features of speech in absence of visual information. For this, we separately predicted horizontal and vertical eye movements from the acoustic speech envelope using temporal response functions (TRFs). The resulting model fit (i.e. correlation between true and predicted eye movements) is commonly referred to as “speech tracking”. Bayesian regression models were applied to evaluate tracking effects under different conditions of selective attention (single speaker, attended multi-speaker, unattended multi-speaker). Furthermore, we assessed whether individual prediction tendency or semantic word surprisal influenced ocular speech tracking. For vertical eye movements, we found evidence for attended speech tracking in a single speaker condition (*β* = 0.012, 94%HDI = [0.001, 0.0023]) but not in a multi-speaker condition (*β* = 0.006, 94%HDI = [-0.005, 0.016]; see Figure 4A). There was no evidence for tracking of the distracting speech stream (*β* = -0.008, 94%HDI = [-0.020, 0.003]). On the contrary, horizontally directed eye movements selectively track attended (*β* = 0.014, 94%HDI = [0.005, 0.024]), but not unattended (*β* = -0.002, 94%HDI = [-0.011, 0.007]) acoustic speech in a multi-speaker condition (see Figure 4B). Speech tracking in a single speaker condition did not reach significance (*β* = 0.009, 94%HDI = [-0.001, 0.017]) for horizontal eye movements. These findings indicate that eye movements selectively track attended, but not unattended acoustic speech. Furthermore, there seems to be a dissociation between horizontal and vertical ocular speech tracking, indicating that horizontal movements track attended speech in a multi-speaker condition, whereas vertical movements track attended speech in a single speaker condition (see Table 2 for a summary of ocular speech tracking effects).

**Figure 4:**
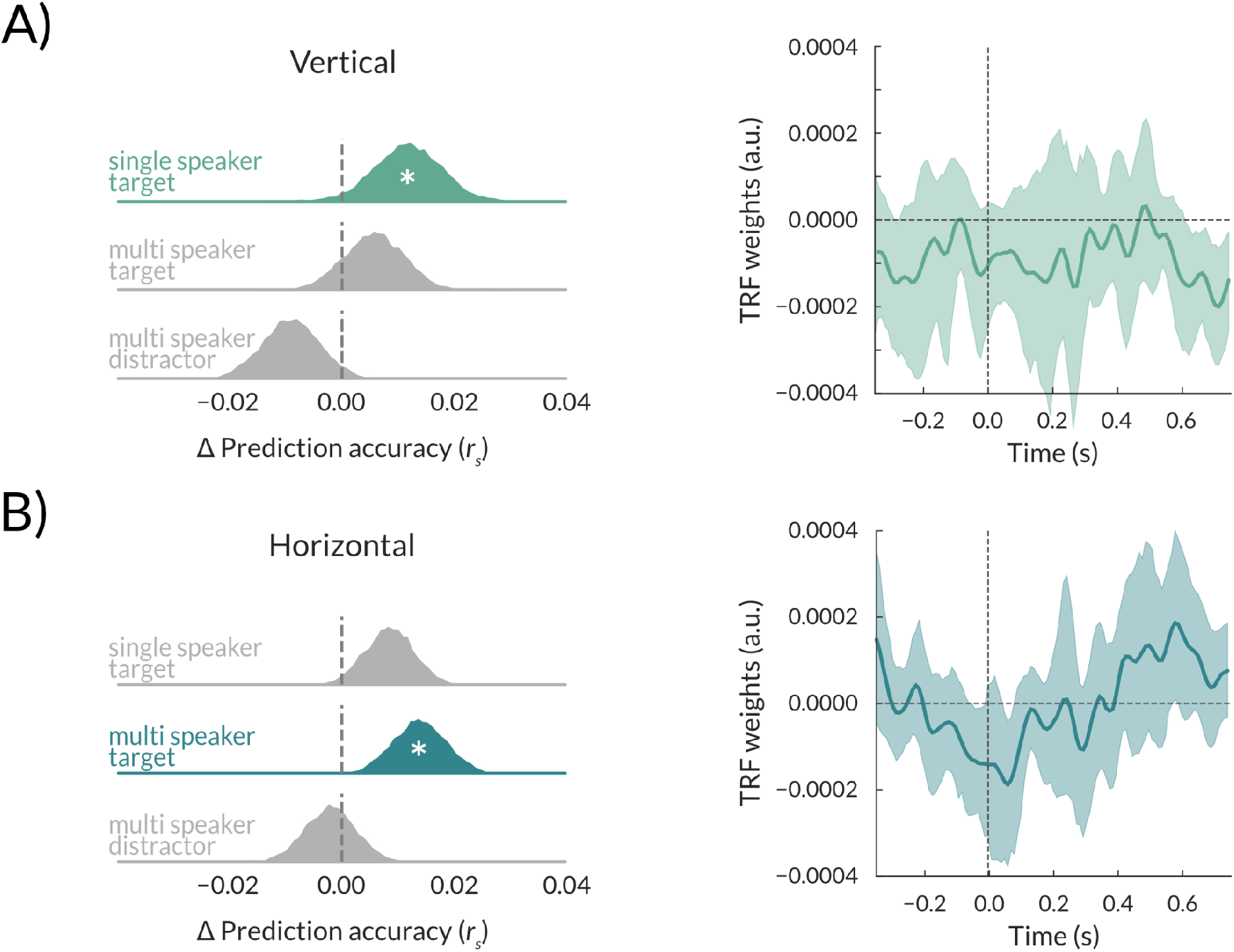
*Ocular speech tracking is dependent on selective attention.* **A)** Vertical eye movements ‘significantly’ track attended clear speech, but not in a multi-speaker condition. Temporal profiles of this effect show a downward pattern (negative TRF weights). **B)** Horizontal eye movements ‘significantly’ track attended speech in a multi-speaker condition. Temporal profiles of this effect show a left-rightwards (negative to positive TRF weights) pattern. Statistics were performed using Bayesian regression models. A ‘*’ within posterior distributions depicts a significant difference from zero (i.e. the 94%HDI does not include zero). Shaded areas in TRF weights represent 95% confidence intervals. N= 29

**Table 2:** Model summary statistics for ocular speech tracking depending on condition and prediction tendency Note: Dependent Variable = ocular speech tracking (correlation between true and predicted eye movements).

|  | vertical eye movements |  |  |  | horizontal eye movements |  |  |  |
| --- | --- | --- | --- | --- | --- | --- | --- | --- |
|  | b | sd | hdi<br>3% | hdi<br>97% | b | sd | hdi<br>3% | hdi<br>97% |
| cond1: single speaker - attended | 0.012 | 0.006 | 0.001 | 0.023 | 0.009 | 0.005 | -0.001 | 0.017 |
| cond2: multi-speaker - attended | 0.006 | 0.006 | -0.005 | 0.016 | 0.014 | 0.005 | 0.005 | 0.024 |
| cond3: multi-speaker - unattended | -0.008 | 0.006 | -0.020 | 0.003 | -0.002 | 0.005 | -0.011 | 0.007 |
| prediction tendency | 0.001 | 0.004 | -0.005 | 0.008 | -0.001 | 0.003 | -0.007 | 0.004 |

Additionally, we wanted to investigate if predictive processes are related to ocular speech tracking using the same approach as for neural speech tracking (see previous section). Crucially, we found no evidence for a relationship between individual prediction tendency and ocular speech tracking on a vertical (*β* = 0.001, 94%HDI = [-0.005, 0.008]) or horizontal (*β* = -0.001, 94%HDI = [-0.007, 0.004]) plane. Similarly, we found no difference in ocular speech tracking between words of high surprisal and their lexically matched controls (see **Table 3**). These findings indicate that individuals engage in ocular speech tracking independent of their individual prediction tendency or overall semantic probability.

**Table 3:** Model summary statistics for ocular speech tracking depending on word type and condition. Note: Dependent Variable = ocular speech tracking (correlation between true and predicted eye movements * Intercept represents speech tracking for lexical control words in a single speaker condition.).

|  | vertical eye movements |  |  |  | horizontal eye movements |  |  |  |
| --- | --- | --- | --- | --- | --- | --- | --- | --- |
|  | b | sd | hdi<br>3% | hdi<br>97% | b | sd | hdi<br>3% | hdi<br>97% |
| Intercept* | 0.053 | 0.019 | 0.019 | 0.092 | 0.026 | 0.018 | -0.008 | 0.061 |
| wordtype (semantic violation effect) | -0.024 | 0.026 | -0.074 | 0.023 | 0.028 | 0.025 | -0.019 | 0.075 |
| condition (multi-speaker effect) | -0.005 | 0.025 | -0.052 | 0.042 | -0.007 | 0.025 | -0.053 | 0.043 |
| wordtype x condition | 0.008 | 0.036 | -0.058 | 0.077 | 0.001 | 0.035 | -0.065 | 0.068 |

### Neural speech tracking is mediated by eye movements

Additionally, we performed a similar, but more detailed mediation analysis compared to Gehmacher and colleagues (2024) in order to separately investigate the contribution of horizontal and vertical eye movements to neural speech tracking across different time lags. Following mediation analysis requirements, only significant ocular speech tracking effects were further considered, i.e. vertical eye movements in the clear speech condition and horizontal eye movements in response to a target in the multi-speaker condition. We compared the plain effect (c) of neural speech tracking (using a simple stimulus model with speech envelope and a circularly time-shifted version of the respective eye movements as the predictors for neural responses) to its direct (residual) effect (c’) by including true horizontal or vertical eye movements as a second predictor into the stimulus model. The decrease in predictor weights from the plain to the residual stimulus model indicates the extent of the mediation effect. This model evaluates to what extent gaze behaviour functions as a mediator between acoustic speech input and brain activity. To establish a time-resolved mediation analysis in source-space, we computed the first three principal components (PCs) of the effect (via PCA, see Figure 5).

**Figure 5.**
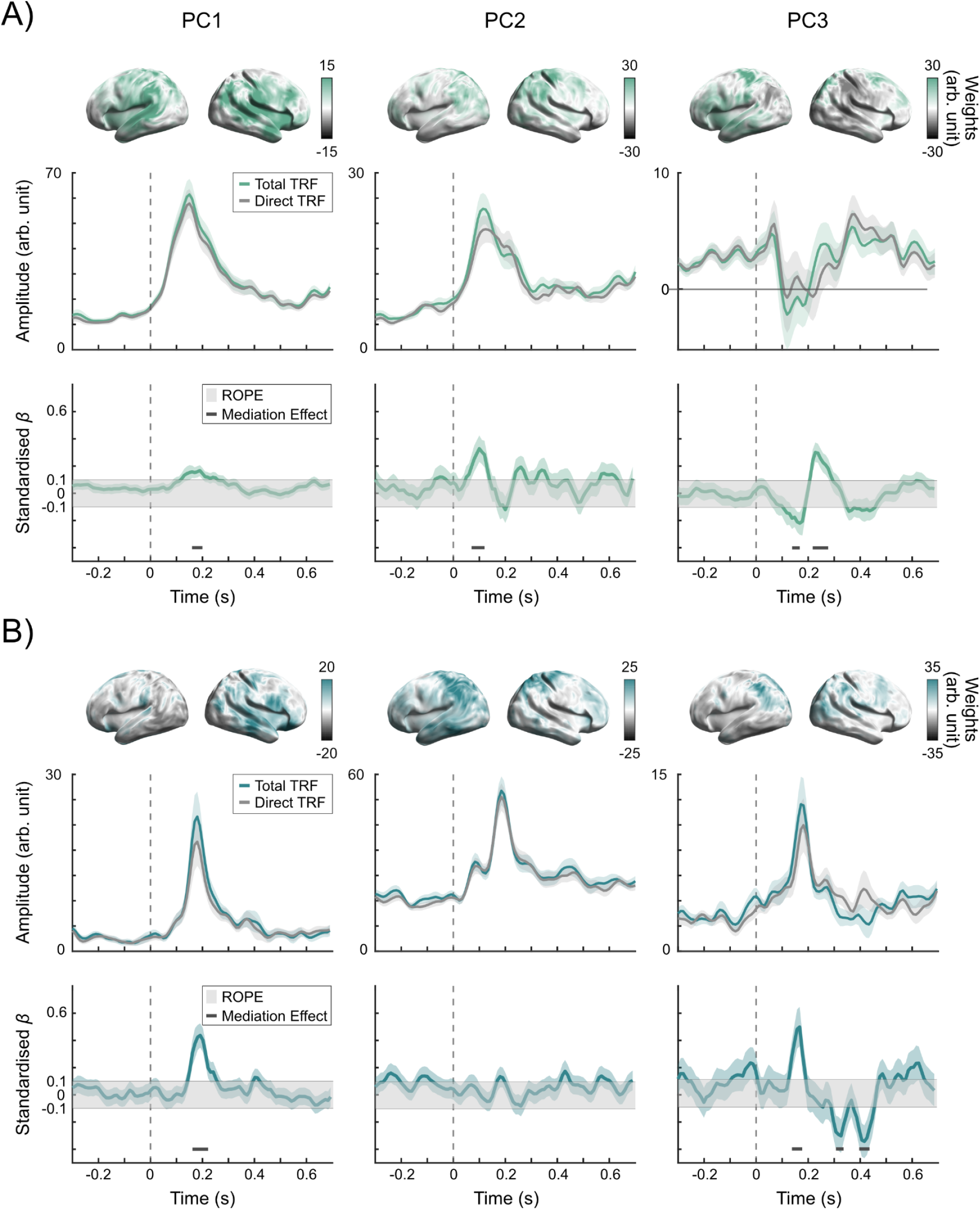
Ocular speech tracking and selective attention to speech share underlying neural computations. (**A**) Vertical eye movements mediate neural tracking of clear speech. *Top:* Spatial distribution (PCA weights) of the group-averaged mediation effect for the first three principal components (PC1–PC3). Mediation effects are distributed across bilateral language networks, including early sensory, parietal, frontal, and lateral semantic processing areas. *Middle:* Amplitude of the Total Temporal Response Function (TRF; green) versus the Direct TRF (grey) over time. *Bottom:* Time-resolved standardized beta coefficients representing the mediation effect. Vertical eye movements significantly mediate neural tracking at early latencies (0.07–0.28 s). (**B**) Horizontal eye movements contribute to the neural tracking of target speech in a multi-speaker environment. Layout is identical to (A), with Total TRF shown in blue. Horizontal eye movements show similar early mediation effects alongside two additional negative clusters, indicating later effects at ∼300 ms and ∼400 ms respectively. Shaded ribbons around the waveform lines indicate mean and standard error for the TRF comparison panels and the mean and 94% HDI for the mediation effect panels. In the bottom panels, the grey shaded rectangular background band represents the Region of Practical Equivalence (ROPE; Kruschke, 2018). Solid horizontal grey markers denote robust clusters where a minimum of two contiguous time-points exhibited a significant mediation effect. Statistical evaluations were performed using Bayesian regression models (N = 29).

We observed significant mediation for all three PCs during vertical eye movements in clear speech (Figure 5A), but only for PC1 and PC3 during horizontal eye movements in the multi-speaker condition (Figure 5B). PC1 exhibited a robust, sustained mediation effect (-0.3 to 0.7 s) peaking at ∼0.18 s across widespread auditory regions, with a right-hemispheric dominance in both conditions. PC2 loaded predominantly on left auditory regions with an early peak (∼0.08 s). While this effect was significant for vertical movements in clear speech, the corresponding left-parietal component for horizontal movements in the multi-speaker condition did not reach group-level significance, reflecting high inter-subject variability within this specific network. PC3 revealed a brief negative cluster followed by a positive one indicating a latency shift (i.e. delayed model weights for neural responses when oculomotor responses are included) for vertical movements. For horizontal movements, PC3 maintained robust effects in left auditory areas with discrete, short-latency peaks (∼ -0.2 s, ∼0.05 s). Notably, horizontal movements also yielded late negative clusters (∼0.3 s and ∼0.4 s). In this context, negative clusters could indicate reversed effects (i.e. model weights for neural responses are increased when oculomotor responses are included), which are difficult to interpret as they allow for several explanations (such as anticorrelated data, larger shifts in latency or changes in temporal precision). Taken together, this suggests that eye movements contribute considerably to neural speech tracking over widespread cortical areas that are commonly related to speech processing and attention.

### Neural and ocular speech tracking are differently related to comprehension

In a final step, we addressed the behavioural relevance of neural as well as ocular speech tracking respectively. At the end of every trial, participants were asked to evaluate 4 different true or false statements about the target story. The accuracy of these responses was averaged within condition (single vs. multi-speaker) and served as an approximation for semantic speech comprehension. Additionally, we evaluated the averaged subjective ratings of difficulty (which were given on a 5-point likert scale).

To avoid calculating separate models for all 102 magnetometers, neural encoding was averaged over selected channels (10% showing the strongest encoding effect). Bayesian regression models were used to investigate relationships between neural/ocular speech tracking and comprehension or difficulty. Ocular speech tracking was analyzed separately for horizontal and vertical eye movements. We found no significant relationship between neural speech tracking and comprehension (*β* = 0.138, 94%HDI = [-0.050, 0.330]), no interaction between neural speech tracking and condition (*β* = -0.088, 94%HDI = [-0.390, 0.201]), but a significant effect for condition (*β* = -0.438, 94%HDI = [-0.714, -0.178]), indicating that comprehension was decreased in the multi-speaker condition (see Figure 6A). Similarly, we found no effect of prediction tendency on comprehension (*β* = 0.050, 94%HDI = [-0.092, 0.197]). When investigating subjective ratings of task difficulty, we also found no effect for neural speech tracking (*β* = 0.156, 94%HDI = [-0.079, 0.382]), no interaction between neural speech tracking and condition (*β* = 0.041, 94%HDI = [-0.250, 0.320]), nor individual prediction tendency (*β* = -0.058, 94%HDI = [-0.236, 0.127]). There was, however, a significant difference between conditions (*β* = 1.365, 94%HDI = [1.128, 1.612]), indicating that the multi-speaker condition was rated more difficult than the single speaker condition. In contrast to neural findings, we found a negative relationship between ocular speech tracking and comprehension for vertical as well as horizontal eye movements irrespective of condition (see Figure 6B-C and Table 4). This suggests that participants with weaker performance in semantic comprehension increasingly engaged in ocular speech tracking. There was, however, no significant relationship between subjectively rated difficulty and ocular speech tracking (see Table 5). Presumably, subjective ratings are less comparable between participants, which might be one reason why we did find an effect for objective but not subjective measures.

**Figure 6:**
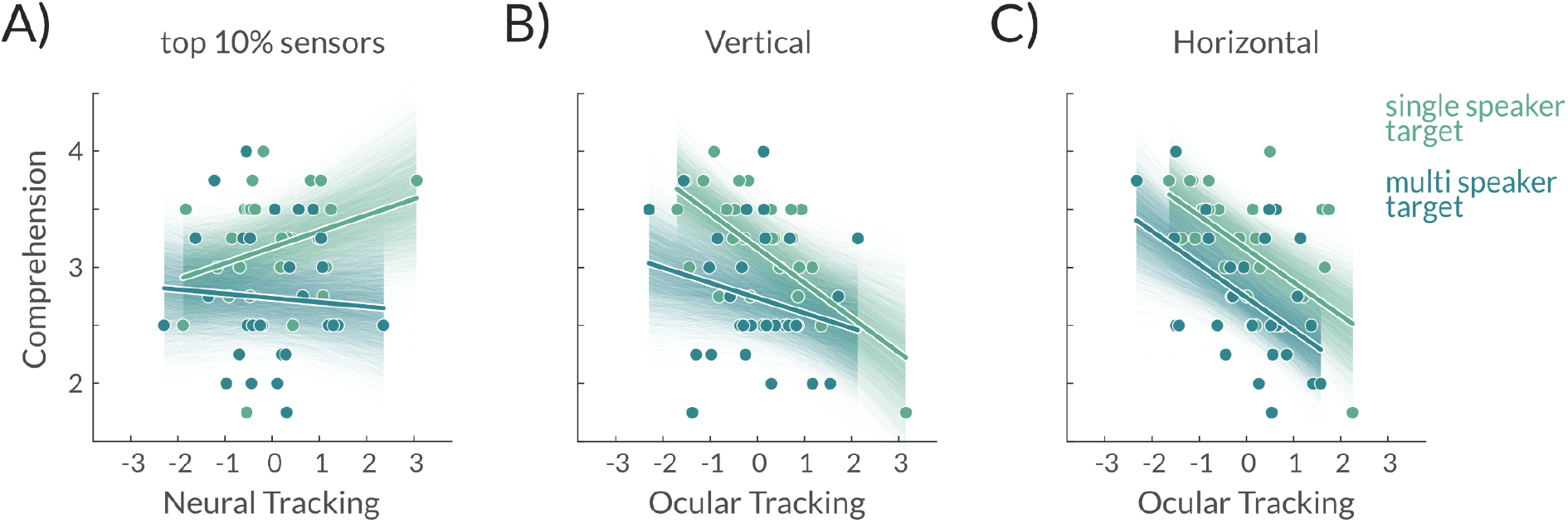
*Ocular, but not neural speech tracking is related to semantic speech comprehension.* **A)** There was no significant relationship between neural speech tracking (10% sensors with strongest encoding effect) and comprehension, however, a condition effect indicated that comprehension was generally decreased in the multi-speaker condition. **B & C)** A ‘significant’ negative relationship between comprehension and vertical as well as horizontal ocular speech tracking shows that participants with weaker comprehension increasingly engaged in ocular speech tracking. Statistics were performed using Bayesian regression models. Shaded areas represent 94% HDIs. N = 29

**Table 4:** Model summary statistics for comprehension depending on ocular speech tracking and condition Note: Dependent Variable = comprehension (averaged accuracy for comprehension questions). * Intercept represents comprehension for average speech tracking in a single speaker condition.

|  | vertical eye movements |  |  |  | horizontal eye movements |  |  |  |
| --- | --- | --- | --- | --- | --- | --- | --- | --- |
|  | b | sd | hdi<br>3% | hdi<br>97% | b | sd | hdi<br>3% | hdi<br>97% |
| Intercept* | 3.17 | 0.099 | 2.986 | 3.356 | 3.162 | 0.095 | 2.972 | 3.333 |
| ocular speech tracking | -0.304 | 0.099 | -0.491 | -0.121 | -0.293 | 0.101 | -0.489 | -0.104 |
| condition (multi-speaker effect) | -0.431 | 0.132 | -0.683 | -0.195 | -0.419 | 0.125 | -0.655 | -0.180 |
| speech tracking x condition | 0.181 | 0.140 | -0.074 | 0.456 | 0.010 | 0.134 | -0.230 | 0.277 |

**Table 5:** Model summary statistics for rated difficulty depending on ocular speech tracking and condition. Note: Dependent Variable = rated difficulty (averaged subjective ratings on a 5-point likert scale). * Intercept represents rated difficulty for average speech tracking in a single speaker condition.

|  | vertical eye movements |  |  |  | horizontal eye movements |  |  |  |
| --- | --- | --- | --- | --- | --- | --- | --- | --- |
|  | b | sd | hdi<br>3% | hdi<br>97% | b | sd | hdi<br>3% | hdi<br>97% |
| Intercept* | 2.186 | 0.123 | 1.971 | 2.429 | 2.192 | 0.122 | 1.971 | 2.430 |
| ocular speech tracking | 0.099 | 0.106 | -0.094 | 0.304 | 0.101 | 0.116 | -0.115 | 0.317 |
| condition (multi-speaker effect) | 1.356 | 0.127 | 1.102 | 1.583 | 1.355 | 0.135 | 1.095 | 1.606 |
| speech tracking x condition | -0.077 | 0.144 | -0.342 | 0.194 | -0.099 | 0.157 | -0.406 | 0.185 |

In sum, the current findings show that ocular speech tracking is differently related to comprehension than neural speech tracking, suggesting that they might not refer to the same underlying concept (e.g. improved representations vs. increased attention). A mediation analysis, however, suggests that they are related and that ocular speech tracking contributes to neural speech tracking (or vice versa).

## DISCUSSION

In the current study, we aimed to replicate and extend findings from two recent studies from our lab in order to integrate them into a comprehensive view of speech processing. In Schubert and colleagues (2023), we found that individual prediction tendencies are related to cortical speech tracking. In Gehmacher and colleagues (2024), we found that eye movements track acoustic speech in selective attention (a phenomenon that we termed “ocular speech tracking”). In the present study, we were able to replicate both findings, providing further details and insight into these phenomena.

In a first step, we confirmed that individuals with a stronger prediction tendency (which was inferred from anticipatory probabilistic sound representations in an independent paradigm) showed an increased neural speech tracking over left frontal areas. Thus, the finding that prediction tendencies generalise across different listening situations seems to be robust. It is assumed that predictions are directly linked to the interpretation of sensory information. This interpretation a) is likely to occur simultaneously at different places in the cognitive (and anatomical) hierarchy and b) describes a bidirectional process in which internal models influence and are influenced by sensory input (Knill & Pouget, 2004). It should be noted that our measure of individual prediction tendency makes no claim on the relative weighting of top-down or bottom-up processing; it does, however, infer an absolute tendency to generate anticipatory (top-down) predictions. This type of prediction is relevant for acoustic processing such as speech and music, whose predictability unfolds over time.

The replicable association between individual differences in this prediction tendency and speech processing stresses the importance of research focusing on the beneficial aspects of individual “traits” in predictive processing. As suggested in a comparative study (Schubert et al., 2024), these predictive tendencies or traits do not generalise across modalities but seem to be reliable within the auditory modality. We suggest that they could serve as an independent predictor of linguistic abilities, complementing previous research on statistical learning (e.g. Siegelman & Frost, 2015).

In a second step, we were able to support the assumption that eye movements track prioritised acoustic modulations in a continuous speech design without visual input. With the current (extended) conceptual replication, we were able to address an important consideration from Gemacher and colleagues (2024): Ocular speech tracking is not restricted to a simplistic 5-word sentence structure but can also be found for continuous, narrative speech. Importantly, in contrast to Gehmacher et al. (2024), we did not observe ocular tracking of the multi-speaker distractor in this study. This difference is likely attributable to the simplistic single-trial, 5-word task structure in Gehmacher et al., which resulted in high temporal overlap between the target and distractor speech streams and likely drove the significant distractor-tracking effects observed in that study. The absence of such an effect during continuous listening in our study suggests that ocular tracking is indeed more specific to selective attention. However, in line with previous results, we found that horizontal ocular speech tracking is increased in a multi-speaker (compared to a single speaker) condition in response to an attended target but not a distractor speaker. In contrast, we found that vertical eye movements solely track auditory information in a single speaker condition. The differential contribution of vertical vs. horizontal eye movements to auditory processing is not a widely studied topic in neuroscience and further replications are necessary to establish the robustness of this dissociation. One possible explanation for these findings is, that in the multi-speaker condition, the demand for spatial segregation was increased compared to the single speaker condition. Although speech was always presented at phantom centre even for both speakers during the multi-speaker condition, multiple speakers are distributed mostly horizontally in natural human environments (e.g. the classic cocktailparty situation; Cherry, 1953). The observed effect might resemble a residual of this learned association between a spatially segregated target speaker’s acoustic output and a lateral shift of gaze (and attention) on the horizontal plane towards the target’s source location, maximising information and thereby minimising uncertainty about the targeted auditory object. Relatedly, it remains an open question whether microsaccades are a key feature driving ocular speech tracking. However, our current study does not analyze microsaccades due to methodological constraints: microsaccades are binary response vectors, which are incompatible with TRF analyses used here. Addressing this would require adapting models to handle time-continuous binary response data or potentially exploring alternative approaches, such as regression-based ERFs (e.g., as in Heilbron et al., 2022). While these limitations preclude microsaccade analysis in the current study, we hypothesize that they could enhance temporal precision and selectively amplify relevant sensory input, supporting auditory perception. Future studies should explore this possibility to uncover the specific contributions of microsaccades to speech tracking.

Irrespective of the potential differences in gaze direction and condition, we found that ocular movements contribute to neural speech tracking over widespread language processing networks, replicating and extending the sensor-level findings of Gehmacher et al. (2024) with time-resolved analyses of this mediation effect. It is important to note that our current findings do not allow for inference on directionality. Our choice of ocular movements as a mediator was motivated by the fact that the relationship between acoustic speech and neural activity is well established, as well as previous results indicating that oculomotor activity contributes to cognitive effects in auditory attention (Popov et al., 2022). However, an alternative model may suggest that neural activity mediates the effect of ocular speech tracking. Hence, it is possible that ocular mediation of speech tracking may reflect a) active (ocular) sensing for information driven by (top-down) selective attention or b) improved neural representations as a consequence of temporally aligned increase of sensory gain or c) (not unlikely) both. In fact, when rejecting the notion of a single bottom-up flow of information and replacing it with a model of distributed parallel and dynamic processing, it seems only reasonable to assume that the direction of communication (between our eyes and our brain) will depend on where (within the brain) as well as when we look at the effect. Thus, the regions and time-windows reported here should be taken as an illustration of oculo-neural communication during speech processing rather than an attempt to "explain" neural speech processing by ocular movements.

Despite the finding that eye movements mediate neural speech tracking, the behavioural relevance for semantic comprehension appears to differ between ocular and neural speech tracking. Specifically, we found a negative association between ocular speech tracking and comprehension, indicating that participants with lower comprehension performance exhibited increased ocular speech tracking. Interestingly, no significant relationship was observed between neural tracking and comprehension.

In this context, the negative association between ocular tracking and comprehension might reflect individual differences in how participants allocate cognitive resources. Participants with lower comprehension may rely more heavily on attentional mechanisms to process acoustic features, as evidenced by increased ocular tracking. This reliance could represent a compensatory strategy when higher-order processes, such as semantic integration or memory retrieval, are less effective. Importantly, our comprehension questions (see Experimental Procedure) targeted a broad range of processes, including intelligibility and memory, suggesting that this relationship reflects a trade-off in resource allocation between low-level acoustic focus and integrative cognitive tasks.

Rather than separating eye and brain responses conceptually, our analysis highlights their complementary contributions. Eye movements may enhance neural processing by increasing sensitivity to acoustic properties of speech, while neural activity builds on this foundation to integrate information and support comprehension. Together, these systems form an interdependent mechanism, with eye and brain responses working in tandem to facilitate different aspects of speech processing.

This interpretation is consistent with the absence of a difference in ocular tracking for semantic violations (e.g., words with high surprisal versus lexically matched controls), reinforcing the view that ocular tracking primarily reflects attentional engagement with acoustic features rather than direct involvement in semantic processing. This aligns with previous findings that attention modulates auditory responses to acoustic features (e.g., Forte et al., 2017), further supporting the idea that ocular tracking reflects mechanisms of selective attention rather than representations of linguistic content.

Future research should investigate how these systems interact and explore how ocular tracking mediates neural responses to linguistic features, such as lexical or semantic processing, to better understand their joint contributions to comprehension.

Interestingly, we were not able to relate individual prediction tendency to ocular speech tracking. This raises the question whether ocular speech tracking reflects attentional mechanisms rather than predictive processes. Indeed, the current findings suggest that prediction tendencies and active ocular sensing are related to different aspects of neural speech processing. We propose a perspective in which active sensing is motivated by selective attention, whereas anticipatory prediction tendency is an independent correlate of neural speech tracking. Even though predictive processing as well as attention might be considered as the two pillars on which perception rests, research investigating their selective contributions and interactions is rare. Summerfield and de Lange (2014) have argued that predictions and attention are distinctive mechanisms as the former refers to the probability and the latter to the relevance of an event, arguing that events can be conditionally probable but irrelevant for current goals and vice versa. Similarly, it has been proposed that attention can be integrated into the predictive coding framework in reference to the optimization of sensory precision (Feldman & Friston, 2010). In this view, predictions encode probabilistic representations of a feature and “prediction tendency” refers to the individual reliance on these representations, whereas selective attention determines the precision of sensory inflow that leads to internal model updating. This interpretation is in line with our finding that a) anticipatory feature-specific predictions can be found in a passive listening task, b) prediction tendencies are not increasingly linked to speech tracking with increasing demands on selective attention, whereas on the other hand c) ocular speech tracking seems to increase with selective attention (at least on a horizontal plane), d) remains unaffected by semantic probability and e) contributes to neural speech processing over widespread auditory and parietal areas with a potential overlap with attentional networks. For this reason, we refer to ocular speech tracking as an “active sensing” mechanism that implements the attentional optimization of sensory precision. Instead of passively transducing any input into neural activity, we actively engage with our environment to maximise information, i.e. reduce uncertainty. It has been suggested that motor routines contribute to temporal precision of selective attention, largely determining sensory inflow and hence perception (Morillon et al., 2015; Schroeder et al., 2010). In the current framework, we refer to active sensing via eye movements as shifts of gaze at exact points in time aligned with intensity (information content) fluctuations. In particular, active ocular sensing may even help to increase sensory gain already in the auditory periphery at specific intervals, synchronised with the temporal modulation of attended (but not unattended) speech. As a consequence of this “sensory gating” already at early stages, eye movements potentially affect auditory processing from the ear to the cortex (also see Leszczynski et al., 2023 for saccadic modulations of cortical excitability in auditory areas), contributing to neural speech representations rather than encoding them. Again, this idea is supported by our finding that ocular speech tracking seems to be unaffected by semantic violations.

Indeed, similar active ocular sensing mechanisms have already been suggested to facilitate sound localization (Lovich et al., 2022). Since the current paradigm did not allow for spatial segregation (as competing speech streams have all been presented at phantom centre), it required a different strategy such as (spectro-)temporal differentiation. We argue that active ocular sensing increases temporal precision of complex acoustic input (in order to parse speech into its components of rhythmic, temporally predictable patterns within a particular speaker). Based on the joint findings of the present as well as its preceding studies (Gehmacher et al., 2024, Schubert et al., 2023), we propose a unified working model in which anticipatory predictions as well as active ocular sensing work (independently) together to support auditory speech perception (see Figure 7 for a schematic illustration). We suggest that anticipatory predictions about a feature help to interpret auditory information by prioritizing representational content of high probability at different levels along the perceptual hierarchy. Accordingly, these predictions are formed in parallel and carry high feature-specificity but low temporal precision (as they are anticipatory in nature). This idea is supported by our finding that pure-tone anticipation is visible over a widespread prestimulus interval, instead of being locked to sound onset. It should be noted that the representational content is likely to be different at different levels of the perceptual hierarchy (e.g. encoding of phonemes, words, semantics or even more abstract speaker intentions). Even though the terminology is suggestive of a fixed sequence (similar to a multi storey building) with levels that must be traversed one after each other (and even the more spurious idea of a rooftop, where the final perceptual experience is formed and stored into memory), we distance ourselves from these (possibly unwarranted) ideas. Our usage of “higher” or “lower” simply refers to the observation that the probability of a feature at a higher (as in more associative) level affects the interpretation (and thus the representation and prediction) of a feature at lower (as in more segregated) levels (Caucheteux et al., 2023). Our own findings suggest that individuals differ in their general tendency to create such anticipatory predictions, which leads to differences in neural speech tracking.

**Figure 7:**
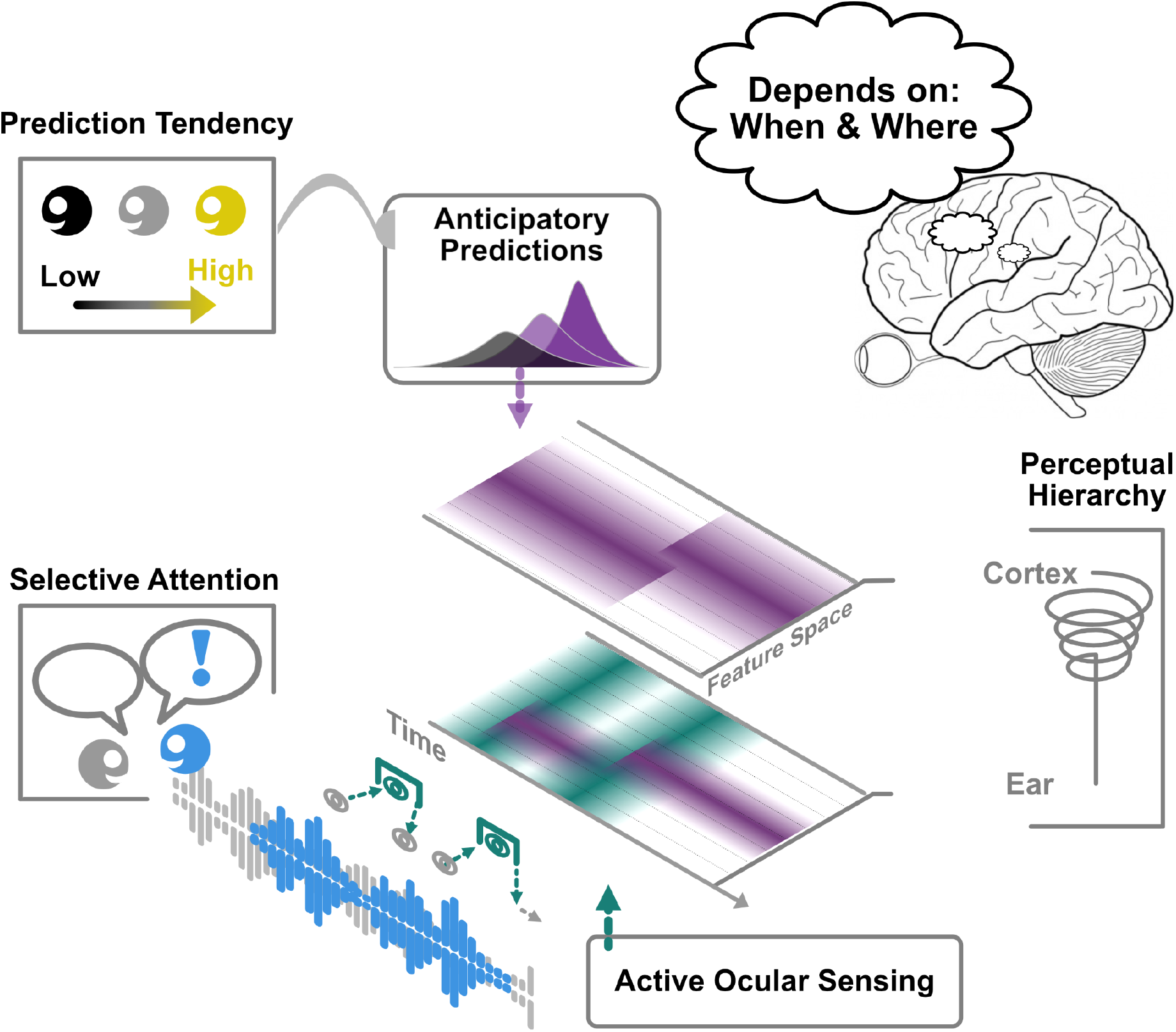
*A schematic illustration of the framework:* A speech percept is processed in representational (feature-)spacetime (X and Y-axes) at different levels of the cognitive hierarchy (Z-axis), which ranges from highly selective to more associative representations (without the notion of an “apex” in this hierarchy). The temporal and spatial characteristics of a representation depends on where and when (in the brain) it is probed. Anticipatory predictions (purple) help to interpret auditory information at different levels in parallel with high feature-specificity but low temporal precision. These anticipatory predictions reflect to some extent individual tendencies (and differences) that generalise across listening situations. In contrast, active ocular sensing (green) increases the temporal precision already at lower stages of the auditory system to facilitate bottom–up processing at specific timescales (similar to neural oscillations). It does not necessarily convey feature-specific information, but is more likely used to boost (or filter) information around relevant time windows. Our results suggest that this mechanism is motivated by selective attention (blue) rather than predictive assumptions

Active ocular sensing, on the other hand, increases temporal precision - potentially already at the early stages of sound transduction - to facilitate bottom-up processing of selectively attended input. Crucially, we suggest that active (ocular) sensing does not necessarily convey feature- or content-specific information, it is merely used to boost (and conversely filter) sensory input at specific timescales (similar to neural oscillations). This assumption is supported by our finding that semantic violations are not differentially encoded in gaze behaviour than lexical controls. Our research suggests that this active ocular sensing mechanism or “filter” is driven by internal (attentional) goals rather than prior beliefs (stressing the distinction to other active sensing mechanisms).

With this speculative framework we attempt to describe and relate three important phenomena with respect to their relevance for speech processing: 1) “Anticipatory predictions” that are created in absence of attentional demands and contain probabilistic information about stimulus features (here, inferred from frequency-specific pre-activations during passive listening to sound sequences). 2) “Selective attention” that allocates resources towards relevant (whilst suppressing distracting) information (which was manipulated by the presence or absence of a distractor speaker). And finally 3) “active ocular sensing”, which refers to gaze behavior that is temporally aligned to attended (but not unattended) acoustic speech input (inferred from the discovered phenomenon of ocular speech tracking). We propose that auditory inflow is, at a basic level, temporally modulated via active ocular sensing, which “opens the gates” in the sensory periphery at relevant timepoints. How exactly this mechanism is guided (for example where the information about crucial timepoints comes from, if not from prediction, and whether it requires habituation to a speechstream etc.) is yet unclear. Unlike predictive tendencies, active ocular sensing appears to reflect selective attention, manifesting as a mechanism that optimizes sensory precision. Individual differences with respect to anticipatory predictions on the other hand, seem to be independent from the other two entities, but nevertheless relevant for speech processing. We therefore support the notion that representational content is interpreted based on prior probabilistic assumptions. If we consider the idea that “a percept” of an (auditory) object is actually temporally and spatially distributed (across representational spacetime - see Fig. 7), the content of information depends on where and when it is probed (see for example Dennett, 1991 for similar ideas on consciousness). Having to select from multiple interpretations across space and time requires a careful balance between the weighting of internal models and the allocation of resources based on current goals. We suggest that in the case of speech processing, this challenge results in an independent adaptation of feature-based precision-weighting by predictions on the one hand and temporal precision-weighting by selective attention on the other.

We suggest that future research on auditory perception should integrate conceptual considerations on predictive processing, active (crossmodal) sensing, and selective attention. In particular, it would be interesting whether ocular speech tracking can be observed for unfamiliar languages with unpredictable prosodic rate. Furthermore, the relationship between neural oscillations in selective attention and active sensing should be further investigated using experimental modulations to address the important, pending question of causality. Brain stimulation (such as tACS; transcranial alternating current stimulation) could be used in an attempt to alter temporal processing frames and / or ocular speech tracking. With regards to the latter, future studies should focus on the potential consequences of inhibited active sensing (e.g. actively disrupting natural gaze behaviour) for neural speech tracking. Our interpretation suggests that increased tracking of unexpected input (i.e. semantic violations) should be affected by active ocular sensing if sensory gain in the periphery is indeed dependent on this mechanism. Currently, the findings propose that active ocular sensing, with its substantial contribution to neural speech tracking, is driven by selective attention, and not by individual differences in prediction tendency.

## ACKNOWLEDGEMENTS

J.S. and Q.G. are supported by the Austrian Science Fund (FWF; Doctoral College "Imaging the Mind"; W 1233-B). Q.G. is also supported by the Austrian Research Promotion Agency (FFG; BRIDGE 1 project “SmartCIs”; 871232) and F.S. is supported by WS Audiology. Thanks to the whole research team. Special thanks to Manfred Seifter for his support in conducting the MEG measurements.

## AUTHOR CONTRIBUTIONS

J.S. and Q.G. designed the experiment, analysed the data, generated the figures, and wrote the manuscript. T.H. recruited participants and supported the data analysis. F.S. supported the data analysis and edited the manuscript. N.W. acquired the funding, supervised the project, and edited the manuscript.

